# To rarefy or not to rarefy: Enhancing diversity analysis of microbial communities through next-generation sequencing and rarefying repeatedly

**DOI:** 10.1101/2020.09.09.290049

**Authors:** Ellen S. Cameron, Philip J. Schmidt, Benjamin J.-M. Tremblay, Monica B. Emelko, Kirsten M. Müller

## Abstract

Amplicon sequencing has revolutionized our ability to study DNA collected from environmental samples by providing a rapid and sensitive technique for microbial community analysis that eliminates the challenges associated with lab cultivation and taxonomic identification through microscopy. In water resources management, it can be especially useful to evaluate ecosystem shifts in response to natural and anthropogenic landscape disturbances to signal potential water quality concerns, such as the detection of toxic cyanobacteria or pathogenic bacteria. Amplicon sequencing data consist of discrete counts of sequence reads, the sum of which is the library size. Groups of samples typically have different library sizes that are not representative of biological variation; library size normalization is required to meaningfully compare diversity between them. Rarefaction is a widely used normalization technique that involves the random subsampling of sequences from the initial sample library to a selected normalized library size. Rarefying is often dismissed as statistically invalid because subsampling effectively discards a portion of the observed sequences. Nonetheless, it remains prevalent in practice. Notably, the superiority of rarefying relative to many other normalization approaches has been argued in diversity analysis. Here, repeated rarefying is proposed as a tool for diversity analyses to normalize library sizes. This enables (i) proportionate representation of all observed sequences and (ii) characterization of the random variation introduced to diversity analyses by rarefying to a smaller library size shared by all samples. While many deterministic data transformations are not tailored to produce equal library sizes, repeatedly rarefying reflects the probabilistic process by which amplicon sequencing data are obtained as a representation of the source microbial community. Specifically, it evaluates which data might have been obtained if a particular sample’s library size had been smaller and allows graphical representation of the effects of this library size normalization process upon diversity analysis results.

## 1. Introduction

Next-generation sequencing (NGS) has revolutionized the understanding of environmental systems through the characterization of microbial communities and their function by examining DNA collected from samples that contain mixed assemblages of organisms (Bartram et al., 2011; Hugerth and Andersson, 2017; Shokralla et al., 2012). It is well known that fewer than 1% of species in the environment can be isolated and cultured, limiting the ability to identify rare and difficult-to-cultivate members of the community (Bodor et al., 2020; Cho and Giovannoni, 2004; Ferguson et al., 1984). In addition to the limitations of culturing, microscopic evaluation of environmental samples remains of limited utility because of challenges in high-resolution taxonomic identification and the inability to infer function from morphology (Hugerth and Andersson, 2017). Metagenomic evaluations employ NGS technology to analyze large quantities of diverse environmental DNA (Thomas et al., 2012) and have largely eliminated challenges associated with culturing and microscopic identification (McMurdie and Holmes, 2014).

Metagenomics encompasses a conglomerate of different sequencing experimental designs, including amplicon sequencing (sequencing of amplified genes of interest) and shotgun sequencing (sequencing of fragments of present genetic material). While shotgun sequencing allows characterization of the entire community, including both taxonomic composition and functional gene profiles, it is not widely accessible due to high sequencing costs and computational requirements for analysis (Bartram et al., 2011; Clooney et al., 2016; Langille et al., 2013). In contrast, the relatively low cost of amplicon sequencing has made it an increasingly popular technique (Clooney et al., 2016; Langille et al., 2013). The amplification and sequencing of specific genes (e.g., taxonomic marker genes) enables characterization of microbial community composition (Hodkinson and Grice, 2015); as a result, it has been successfully applied in many areas of environmental and water research. For example, amplicon sequencing has been used to characterize and predict cyanobacteria blooms (Tromas et al., 2017), describe microbial communities found in aquatic ecosystems (Zhang et al., 2020), and evaluate groundwater vulnerability to pathogen intrusion (Chik et al., 2020). It has also been applied to water quality and treatment performance monitoring in diverse settings (Vierheilig et al., 2015), including drinking water distribution systems (Perrin et al., 2019; Shaw et al., 2015), drinking water biofilters (Kirisits et al., 2019), anaerobic digesters (Lam et al., 2020), and cooling towers (Paranjape et al., 2020).

Processing and analysis of amplicon sequencing data are statistically complicated for a number of reasons (Weiss et al., 2017). In particular, library sizes (i.e., the total number of sequencing reads within a sample) can vary widely among different samples, even within a single sequencing run, and the disparity in library sizes between samples may not represent actual differences in microbial communities (McMurdie and Holmes, 2014). Amplicon sequencing libraries cannot be compared directly for this reason. For example, two replicate samples with 5,000 and 20,000 sequence reads, respectively, are likely to have different read counts for specific sequence variants simply due to the difference in library size. While parametric tools such as generalized linear modelling (e.g., McMurdie and Holmes, 2014) can provide a statistically sound framework for differential abundance analysis, drawing biologically meaningful diversity analysis conclusions from amplicon sequencing data typically requires normalization of library sizes to account for the additional variation in counts that is attributable to differences in library sizes between samples (McKnight et al., 2019). For example, larger samples may appear more diverse than smaller samples (Hughes and Hellmann, 2005). Notably, a variety of normalization techniques that may affect the analysis and interpretation of results have been suggested, including rarefaction (i.e., the process of rarefying libraries to a common size).

Rarefaction is a normalization tool initially developed for ecological diversity analyses to allow for sample comparison without associated bias from differences in sample size (Sanders, 1968). Rarefaction normalizes samples of differing sample size by subsampling each to a shared threshold. Although initially developed for use in ecological studies, rarefaction is a commonly used library size normalization technique for amplicon sequencing data. As a result, it is the subject of considerable debate and statistical criticism (Gloor et al., 2017; McMurdie and Holmes, 2014). Rarefying is typically conducted in a single iteration that only provides a snapshot of the community that might have been observed at the smaller normalized library size. This omits a random subset of observed sequences and potentially also samples with small library sizes and introduces artificial variation to the data (McMurdie and Holmes, 2014).

Repeatedly rarefying, on the other hand, has the potential to address the statistical concerns associated with omission of data and could provide a more statistically acceptable technique than performing a single iteration of rarefying for diversity analyses. It characterizes what data might have been obtained if a particular sample’s library size had been smaller, revealing what can be inferred about community diversity in the source from samples of equal library size. Rarefying repeatedly has received only trivial consideration in the literature (e.g., McMurdie and Holmes, 2014; Navas-Molina et al., 2013).Diversity analysis approaches grounded in statistical inference about source microbial diversity (that address the random probabilistic processes through which NGS yields libraries of sequence reads) could conceptually be superior to rarefying (Willis, 2019), but they are not yet fully developed or readily available for routine diversity analysis to support study of environmental microbial communities.

Here, we investigate the application of repeatedly rarefying as a library size normalization technique specifically for diversity analyses. This paper graphically evaluates the impact of subsampling with or without replacement and normalized library size selection on diversity analyses such as the Shannon index and Bray-Curtis dissimilarity ordinations, specifically. Rather than representing diversity as a single numerical value or point in an ordination plot (often following transformation that may not be designed to compensate for differing library sizes), rarefying repeatedly yields bands of values or patches of points that characterize how diversity may vary among or between samples at a particular library size.

## 2. Theory

### 2.1 Amplicon Sequencing and Diversity Analysis for Microbial Communities in Water – An Overview

Due to the inevitable interdisciplinarity of environmental water quality research and the complexity and novelty of next generation sequencing relative to traditional microbiological methods used in water quality analyses, further detail on amplicon sequencing is provided. Amplification and sequencing of taxonomic marker genes has been used extensively to examine phylogeny, evolution, and taxonomic classification of numerous groups across the three domains of life (Quast et al., 2013; Weisburg et al., 1991; Woese et al., 1990). Taxonomic marker genes include the 16S rRNA gene in mitochondria, chloroplasts, bacteria and archaea (Case et al., 2007; Tsukuda et al., 2017; Weisburg et al., 1991; Yang et al., 2016), or the 18S rRNA gene within the nucleus of eukaryotes (Field et al., 1988). Widely used reference databases have been developed containing marker gene sequences across numerous phyla (Hugerth and Andersson, 2017).

The 16S rRNA gene consists of nine highly conserved regions separated by nine hypervariable regions (V1-V9; Gray et al., 1984) and is approximately 1,540 base pairs in length (Kim et al., 2011; Schloss and Handelsman, 2004). While sequencing of the full 16S rRNA gene provides the highest taxonomic resolution (Johnson et al., 2019), many studies only utilize partial sequences due to limitations in read length of NGS platforms (Kim et al., 2011). Next-generation sequencing on Illumina platforms (Illumina Inc., San Diego, California) produces reads that are up to 350 base pairs in length, requiring selection of an appropriate region of the 16S rRNA gene to amplify and sequence for optimal taxonomic resolution (Bukin et al., 2019; Kim et al., 2011). Sequencing the more conservative regions of the 16S rRNA gene may be limited to resolution of higher levels of taxonomy, while more variable regions can provide higher resolution for the classification of sequences to the genus and species levels in bacteria and archaea (Bukin et al., 2019; Kim et al., 2011; Yang et al., 2016).

Different variable regions of the 16S rRNA gene may be biased towards different taxa (Johnson et al., 2019) and be preferred for different ecosystems (Escapa et al., 2020). For example, the V4 region has been shown to strongly differentiate taxa from the phyla Cyanobacteria, Firmicutes, Fusobacteria, Plantomycetes, and Tenericutes but the V3 region best differentiates taxa from the phyla Proteobacteria (e.g., *Escherichia coli, Salmonella* spp., *Campylobacter* spp.), Acidobacteria, Bacteroidetes, Chloroflexi, Gemmatimonadetes, Nitrospirae, and Spirochaetae (Zhang et al., 2018). The V4 region of the 16S rRNA gene is frequently targeted using specific primers designed to minimize amplification bias while accounting for common aquatic bacteria (Walters et al., 2015) and is frequently used in aquatic studies (Zhang et al., 2018). It is important to consider suitability of a 16S rRNA region for the habitat (Escapa et al., 2020) and the taxa present in the microbial community due to potential bias of analyzing differing subregions of the 16S rRNA gene (Johnson et al., 2019; Zhang et al., 2018).

The use of amplicon sequencing of partial sequences of the 16S rRNA gene allows examination of microbial community composition and the exploration of shifts in community structure in response to environmental conditions (Hodkinson and Grice, 2015), and identification of differentially abundant taxa between samples (Hugerth and Andersson, 2017). Amplicon sequencing datasets can be analyzed using a variety of bioinformatics pipelines for sequence analysis (e.g., sequence denoising, taxonomic classification, diversity analysis) including *mothur* (Schloss et al., 2009) and *QIIME2* (Bolyen et al., 2019). Previously, sequencing analysis involved the creation of dataset-dependent operational taxonomic units (OTUs) by clustering sequences into groups that met a certain similarity threshold, resulting in a loss of representation of variation in sequences and precluding cross-study comparison (Callahan et al., 2017). Advances in computational power have allowed a shift from use of OTUs to amplicon sequence variants (ASVs) representative of each unique sequence in a sample, which allows for the comparison of sequence variants generated in different studies and retains the full observed biological variation (Callahan et al., 2017). The implementation of tools included bioinformatics pipelines, such as *DADA2* (Callahan et al., 2016) or *Deblur* (Amir et al., 2017), allows quality control of sequencing through the removal of sequencing errors and for the creation of ASVs.

Quality controlled sequencing data for a particular run is then organized into large matrices where columns represent experimental samples and rows contain counts for different ASVs (Weiss et al., 2017). Amplicon sequencing samples have a total number of sequencing reads known as the library size (McMurdie and Holmes, 2014), but do not provide information on the absolute abundance of sequence variants (Gloor et al., 2016, 2017). This data can be used for studies on taxonomic composition, differential abundance analysis and diversity analyses (Figure 1). Taxonomic classification of 16S rRNA sequences using rRNA databases including SILVA (Quast et al., 2013), the Ribosomal Database Project (Cole et al., 2014) and GreenGenes (DeSantis et al., 2006) allows for construction of taxonomic community profiles (Bartram et al., 2011). Taxonomic composition analysis allows for characterization of microbial communities by classifying sequence variants based on similarities to sequences in online databases. The creation of taxonomic composition graphs frequently expresses community composition in proportions. Differential abundance analysis is utilized to explore whether specific sequence variants are found in significantly different proportions between samples (Weiss et al., 2017) to identify potential biological drivers for these differences. This application is outside the scope of this work and is frequently performed using programs initially designed for transcriptomics, such as *DESeq2* (Love et al., 2014) and *edgeR* (Robinson et al., 2009), or programs designed to account for the compositional structure of sequence data *ALDeX2* (Fernandes et al., 2014). The final potential application of this data is diversity analyses, which can be evaluated on varying scales from within sample (alpha) to between samples (beta; Sepkoski, 1988) but is associated with the challenge of the true diversity of environmental sources largely remaining unknown (Hughes et al., 2001).

**Figure 1.**
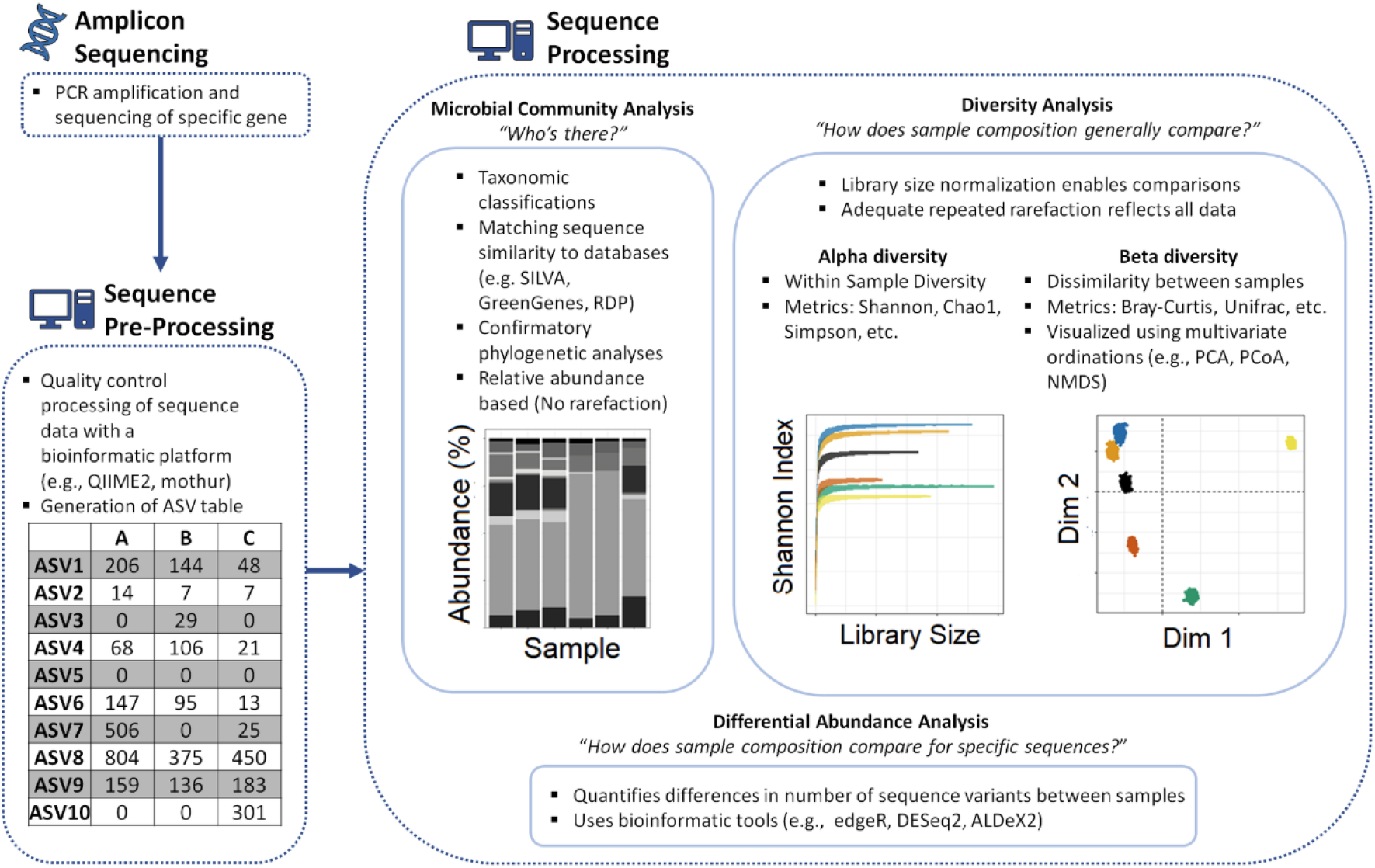
Schematic of general workflow in amplicon sequencing of samples.

Alpha diversity serves to identify richness (e.g., number of observed sequence variants) and evenness (e.g., allocation of read counts across observed sequence variants) within a sample (Willis, 2019). Comparison of alpha diversity among samples of differing library sizes may result in inherent biases, with samples having larger library sizes appearing more diverse due to the potential presence of more sequence variants in samples with larger libraries (Hughes and Hellmann, 2005; Willis, 2019). This has commonly required samples to have equal library sizes before comparison to prevent bias fabricated only from differences in initial library size. Diversity indices used to characterize the alpha diversity of samples include but are not limited to the Shannon index (Shannon, 1948), Chao1 index (Chao and Bunge, 2002), and Simpson index (Simpson, 1949), but unique details of such indices should be understood for correct usage. For example, Chao1 relies on the observation of singletons in data to estimate diversity (Chao and Bunge, 2002), but denoising processes for sequencing data may remove singleton reads making the Chao1 estimator invalid for accurate analysis. The Shannon index, used in this study, is affected by differing library sizes because the contribution of rare sequences to total diversity is progressively lost with smaller library sizes.

Similar to alpha diversity, samples with differing library sizes in beta diversity analyses may produce erroneous results due to the potential for samples with larger library sizes to have more unique sequences simply due to the presence of more sequence variants (Weiss et al., 2017). A variety of beta diversity metrics can be used to compare sequence variant composition between samples including Bray-Curtis (Bray and Curtis, 1957) or Unifrac (Lozupone and Knight, 2007) distances, which can then be visualized using ordination techniques (e.g., PCA, PCoA, NMDS). Bray-Curtis dissimilarity, used in this study, includes pairwise comparison of the numbers for each ASV between two samples, which are expected to be quite dissimilar (even if the communities they represent are not) if library sizes vary substantially.

### 2.2 Limitations of Library Size Normalization Techniques

Diversity analysis, as it is presently applied, usually requires library size normalization to account for bias introduced through varying read counts in samples. For example, samples with larger library sizes may appear more diverse simply due to the presence of more sequences. Normalization techniques that feature various statistical transformations have been proposed for use in place of rarefying or proportions (McKnight et al., 2019), including upper-quartile log fold change (e.g., Robinson et al., 2009), centered log-ratio transformations (e.g., Gloor et al., 2017), geometric mean pairwise ratios (e.g., Chen et al., 2018), variance stabilizing transformations (e.g., Love et al., 2014) or relative log expressions (e.g., Badri et al., 2018). McKnight et al. (2019) noted that the failure of most normalization techniques to transform data to equal library sizes for diversity analysis “is discouraging, as standardizing read depths are the initial impetus for normalizing the data (i.e., if all samples had equal read depths after sequencing, there would be no need to normalize”.

These proposed alternatives to rarefying are also often compromised by the presence of large proportions of zero count data in tabulated amplicon sequencing read counts. Zero counts represent a lack of information (Silverman et al., 2018) and may arise from true absence of the sequence variant in the sample or a loss resulting in it not being detected when it was actually present (Tsilimigras and Fodor, 2016; Wang and LêCao, 2019). Nonetheless, many normalization procedures for amplicon sequencing datasets require zero counts to be omitted or modified, especially when applying transformations that utilize logarithms (e.g., centered log-ratio, relative log expressions, geometric mean pairwise ratios). Methods that utilize logarithms involve fabricating count values (pseudocounts) for the many zeros of which amplicon sequencing datasets are comprised and selecting a pseudocount value is an additional challenge (Weiss et al., 2017) that may be accomplished using probabilistic arguments (Gloor et al., 2016; 2017). Zeros are a natural occurrence in discrete, count-based data such as the counting of microorganisms or amplicon sequences and adjusting or omitting them can introduce substantial bias into microbial analyses (Chik et al., 2018).

McMurdie and Holmes (2014) noted that use of proportions is problematic due to heteroscedasticity: for example, one sequence read in a library size of 100 is a far less precise representation of source composition than 100 sequence reads in a library size of 10,000, even though both comprise 1% of the observed sequences. McKnight et al. (2019) favour use of proportions in diversity analysis without noting how precision of proportions, and the degree to which alpha diversity in the source is reflected (Willis, 2019), varies with library size. Willis (2019) also points towards a conceptually better approach to diversity analysis that accounts for measurement error and the difference between the sample data and the population (environmental source) of which the sample data are only a partial representation. Diversity analysis in general does not do this, as it applies a set of calculations to sample data (or some transformation thereof) to obtain one value of alpha diversity or one point on an ordination plot. Pending further development of such approaches, this study revisits rarefying because of the practical simplicity of comparing diversity among samples of equal library size.

McMurdie and Holmes (2014) propose that rarefying is not a statistically valid normalization technique due to the omission of valid data, which may be resolved for the purposes of diversity analysis by rarefying repeatedly to represent all sequences in the proportions with which they were observed and compare sample-level microbial community diversity at a particular library size. In addition, McMurdie and Holmes (2014) dismissed repeatedly rarefying as a normalization technique, in part because repeatedly rarefying an artificial library consisting of a 50:50 ratio of two sequence variants does not yield a 50:50 ratio at the rarefied library size and this added noise could affect downstream analyses. However, such error is inherent to subsampling, whether from a population or from a larger sequence library and has thus already affected samples with smaller library sizes; it is the reason why simple proportions are less precise in samples with smaller library sizes.

McMurdie and Holmes (2014), also cited the investigation of Navas-Molina et al. (2013) as an example of repeatedly rarefying to normalize library sizes and used it to support their dismissal of this technique due to the omission of valid data and added variability. However, it is critical to note that the work in Navas-Molina et al. (2013) reported using jackknife resampling of sequences, which cannot be equated to repeatedly rarefying (random resampling with or without replacement). Hence, it is necessary to build upon preliminary analysis of repeatedly rarefying as a normalization technique and to explore the impact of subsampling approach and normalized library size on diversity analysis results.

## 3. Methods

### 3.1 Example Data – DNA Extraction and Amplicon Sequencing

Samples used in this study are part of a larger study at Turkey Lakes Watershed (North Part, ON), but only an illustrative subset of samples is considered for the purpose of evaluating rarefaction rather than for ecological interpretation. This allows evaluation of repeated rarefying as a normalization technique without utilizing simulated data. DNA extracts isolated from environmental samples were submitted for amplicon sequencing using the Illumina MiSeq platform (Illumina Inc., San Diego, California) at the commercial laboratory Metagenom Bio Inc. (Waterloo, Ontario). Primers designed to target the 16S rRNA gene V4 region [515FB (GTGYCAGCMGCCGCGGTAA) and 806RB (GGACTACNVGGGTWTCTAAT; Walters et al., 2015)] were used for PCR amplification.

### 3.2 Sequence Processing and Library Size Normalization

The program *QIIME2* (v. 2019.10; Bolyen et al., 2019) was used for bioinformatic processing of sequence reads. Demultiplexed paired-end sequences were trimmed and denoised, including the removal of chimeric sequences and singleton sequence variants to avoid sequences that may not be representative of real organisms, using *DADA2* (Callahan et al., 2016) to construct the ASV table. Zeroing all singleton sequences could erroneously remove legitimate sequences, particularly if the sequence in question is detected in large numbers in other similar samples; however, the potential effect of such error upon diversity analysis is beyond the scope of this work. Output files from *QIIME2* were imported into R (v. 4.0.1; R Core Team, 2020) for community analyses using *qiime2R* (v. 0.99.23; Bisanz, 2018). Initial sequence libraries were further filtered using *phyloseq* (v. 1.32.0; McMurdie and Holmes, 2013) to exclude amplicon sequence variants that were taxonomically classified as mitochondria or chloroplast sequences. We developed a package called *mirlyn* (Multiple Iterations of Rarefaction for Library Normalization; Cameron and Tremblay, 2020) that facilitates implementation of techniques used in this study built from existing R packages (Table S1). Using the output from *phyloseq, mirlyn* was used to (1) generate rarefaction curves, (2) repeatedly rarefy libraries to account for variation in library sizes among samples, and (3) plot diversity metrics given repeated rarefaction.

### 3.3 Community Diversity Analyses on Normalized Libraries

The impact of normalized library size on the Shannon index (Shannon, 1948), an alpha diversity metric, was evaluated. Normalized libraries were also used for beta diversity analysis. A Hellinger transformation was applied to normalized libraries to account for the arch effect regularly observed in ecological count data and Hellinger-transformed data were then used to calculate Bray-Curtis distances (Bray and Curtis, 1957). Principal component analysis (PCA) was conducted on the Bray-Curtis distance matrices.

### 3.4 Study Approach

Typically, rarefaction has only been conducted a single time in microbial community analyses, and this omits a random subset of observed sequences, introducing a possible source of error. To examine this error, samples were repeatedly rarefied 1000 times. This repetition provides a representative suite of rarefied samples capturing the randomness in sequence variant composition imposed by rarefying. The sections below address the various decisions that must be made by the analyst and factors affecting reliability of results when rarefaction is used.

#### 3.4.1 The Effects of Subsampling Approach – With or Without Replacement

Rarefying library sizes may be performed with or without replacement. To evaluate the effects of subsampling replacement approaches, we repeatedly rarefied filtered sequence libraries with and without replacement. Results of the two approaches were contrasted in diversity analyses to evaluate the impact of subsampling approach on interpretation of results.

#### 3.4.2 The Effects of Normalized Library Size Selection

Rarefying involves the selection of an appropriate sampling depth to be shared by each sample. To evaluate the effects of different rarefied library sizes, filtered sequence libraries were rarefied repeatedly to varying depths. Results for various sampling depths were contrasted in diversity analyses to evaluate the impact of normalized library size selection on interpretation of results.

## 4. Results and Discussion

### 4.1 Use of Rarefaction Curves to Explore Suitable Normalized Library Sizes

Rarefying requires the selection of a potentially arbitrary normalized library size, which can impact subsequent community diversity analyses and therefore presents users with the challenge of making an appropriate decision of what size to select (McMurdie and Holmes, 2014). Suitable sampling depths for groups of samples can be determined through the examination of rarefaction curves (Figure 2). By selecting a library size that encompasses the flattening portion of the curve for each sample, it is generally assumed that the normalized library size will adequately capture the diversity within the samples despite the exclusion of sequence reads during the rarefying process (i.e., there are progressively diminishing returns in including more of the observed sequence variants as the rarefaction curve flattens).

**Figure 2.**
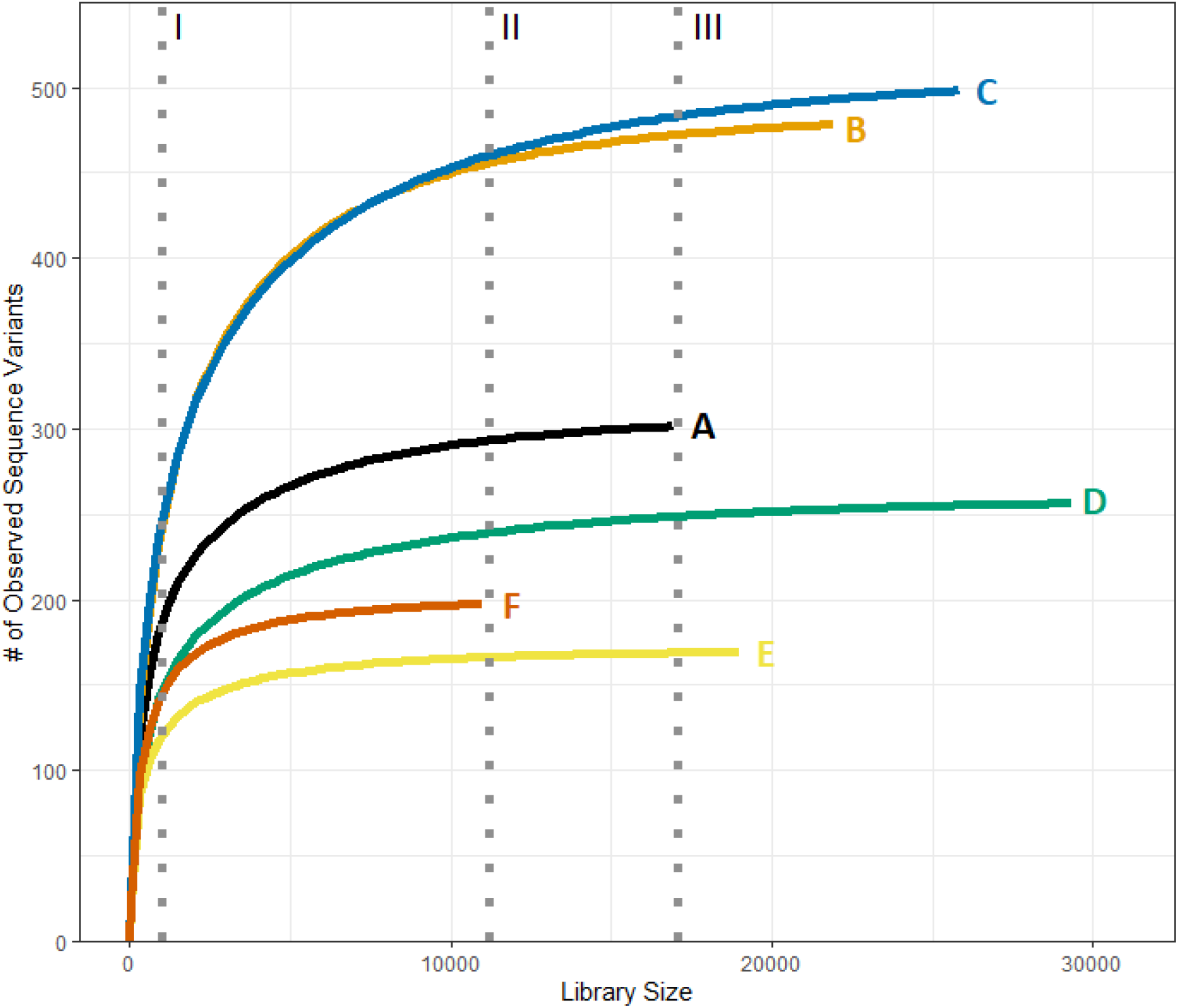
Rarefaction curves showing the number of unique sequence variants as a function of normalized library size for six samples (labelled A – F) of varying diversity and initial library size. Selection of unnecessarily small library sizes (I) omits many sequence variants. Rarefying to the smallest library size (II) omits fewer sequences and variants. While selection of a larger normalized library size (III) would omit even less sequences, it is necessary to omit entire samples (e.g., Sample F) that have too few sequences)

Suggestions have previously been made encouraging selection of a normalized library size that is encompassing of most samples (e.g., 10,000 sequences) and advocation against rarefying below certain depths (e.g., 1,000 sequences) due to decreases in data quality (Navas-Molina et al., 2013). However, generic criteria may not be applicable to all datasets and exploratory data analysis is often required to make informed and appropriate decisions on the selection of a normalized library size. Although previous research advises against rarefying below certain thresholds, users may be presented with the dilemma of selecting a sampling depth that either does not capture the full diversity of a sample depicted in the rarefaction curve (Figure 2 – I) or would require the omission of entire samples with smaller library sizes (Figure 2 – III). The implementation of multiple iterations of rarefying library sizes will aid in alleviating this dilemma by capturing the potential losses in community diversity for samples that are rarefied to lower than ideal depth. Doing so with two or more normalized library sizes may reveal differences in diversity attributable to relatively rare variants that could be suppressed by normalizing to too small of a library size.

### 4.2 The Effects of Subsampling Approach and Normalized Library Size Selection on Alpha Diversity Analyses

The differences in how rarefying samples may be carried out requires users to be diligent in the selection of appropriate tools and commands for their analysis. The R package *phyloseq*, a popular tool for microbiome analyses, has default settings for rarefying including sampling with replacement to optimize computational run time and memory usage (McMurdie and Holmes, 2013). Sampling without replacement, however, is more appropriate statistically because it draws a subset from the observed set of sequences (as though the sample had yielded only the specified library size), whereas sampling with replacement fabricates a set of sequences in similar proportions to the observed set of sequences (Figure 3). Sampling with replacement can potentially cause a rare sequence variant to appear more frequently in the rarefied sample than it was in the original library.

**Figure 3.**
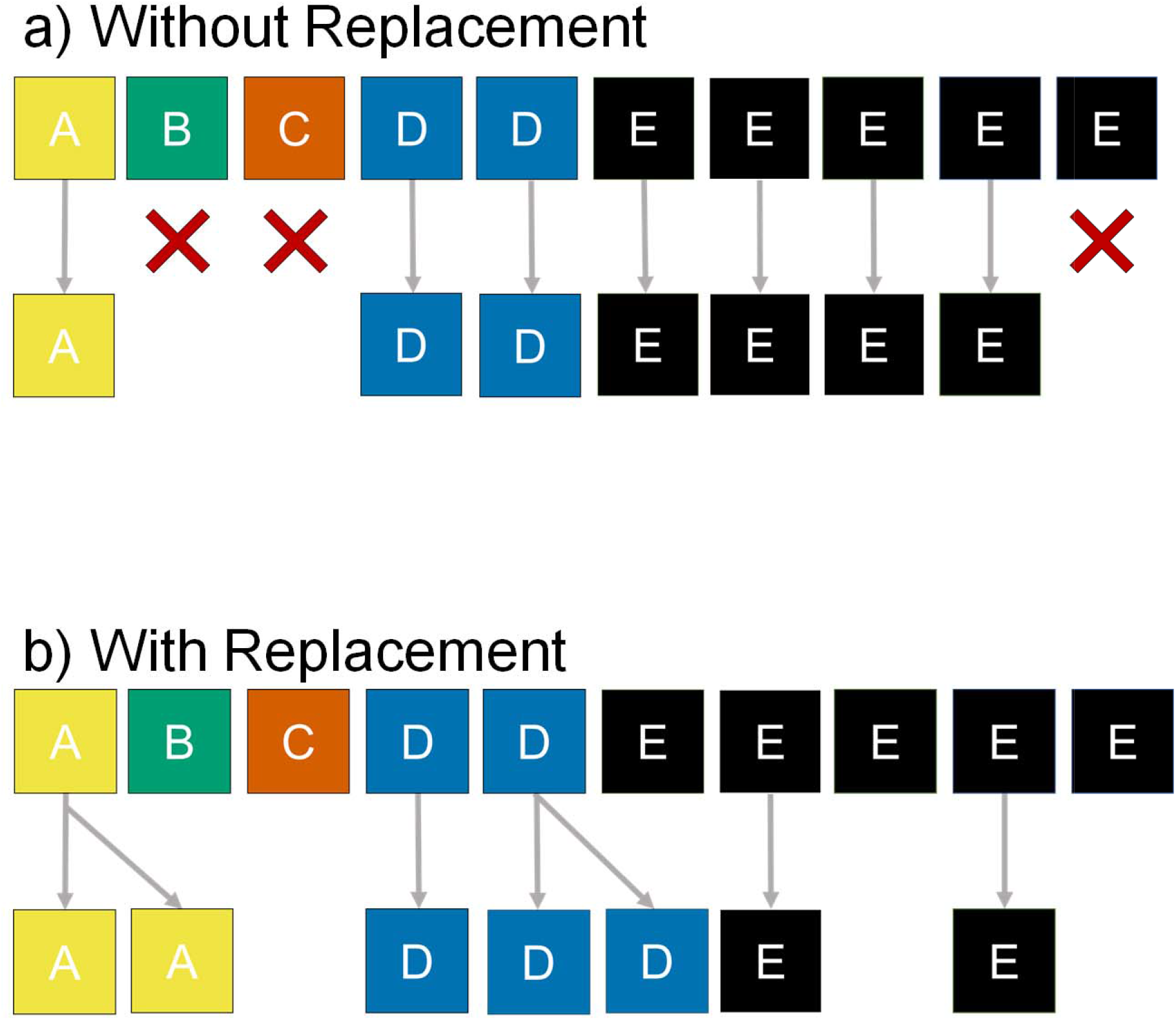
The mechanics of rarefying with or without replacement for a hypothetical sample with a library size of ten composed of five sequence variants (A – E). Rarefying without replacement (a) draws a subset from the observed library excluding the complementary subset, while rarefying with replacement (b) has the potential to artificially inflate the numbers of some sequence variants beyond what was observed.

Rarefying libraries with or without replacement was not found to substantially impact the Shannon index in the scenarios considered in this study (Figure 4-A), but users should still be aware of potential implications of sampling with or without replacement when rarefying libraries. Libraries rarefied with replacement are observed to have a slightly reduced Shannon index relative to libraries rarefied without replacement at many library sizes because rare sequences are excluded more often when sampling with replacement.

**Figure 4.**
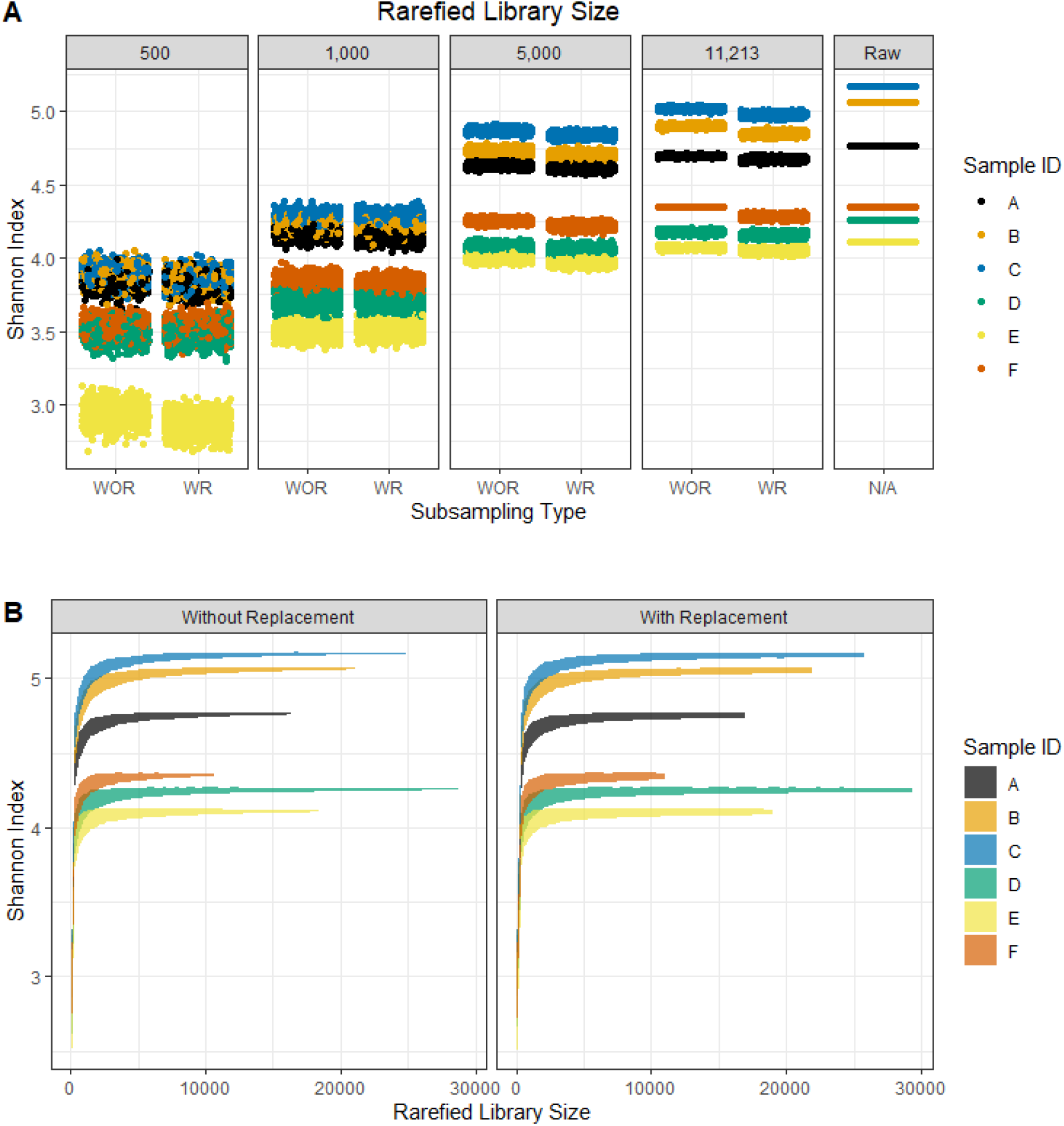
Effect of chosen rarefied library size and sampling with (WR) or without (WOR) replacement upon the Shannon Diversity Index. Six microbial communities were rarefied repeatedly (A) at specific rarefied library sizes of 11,213 sequences, 5,000 sequences, 1,000 sequences, and 500 sequences and (B) to evaluate the Shannon Index as a function of rarefied library size.

The conservation of larger normalized library sizes allows detection of more diversity with minimal variation observed between the iterations of rarefaction (Figure 4-A). The largest considered normalized library size (the sample with the smallest library size has 11,213 sequences) captured the highest Shannon index values, while the Shannon index diminishes for all samples at lower normalized library sizes. The use of repeated iterations of rarefying allows variation introduced through subsampling to be represented in the diversity metric, which is small at larger library sizes. While there was only slight disparity in the Shannon index values between the largest library size and unnormalized data, this may not always be the case and is dependent on the sequence variant composition of the samples. Samples dominated by a large number of low-abundance sequence variants are more likely to have a substantially reduced Shannon index value at a larger normalized library size. Alternatively, samples dominated by only a few highly abundant sequence variants will be comparatively robust to rarefying. A plot of the Shannon index as a function of rarefied library size (Figure 4-B) demonstrates the overall robustness of the Shannon index of these samples for larger library sizes (e.g., > 5,000 sequences) and the increased variation and diminishing values when proceeding to smaller rarefied library sizes. When the normalized library size was decreased to 5,000, the Shannon index is still only slightly reduced by the rarefaction but there is greater variability introduced from rarefying.

The consistency of the diversity metric when rarefying repeatedly is extremely degraded when libraries were rarefied to the smallest considered library size of 500 sequences. It illustrates the potential to reach incorrect conclusions if rarefying is completed only once. When rarefying repeatedly to a small library size, however, diversity index values that are both highly inconsistent and suppressed relative to the diversity of the unrarefied data may lead to inappropriate claims of identical diversity values between samples (e.g., samples A, B, and C become indistinguishable). The extreme reduction and introduced variation of the Shannon index suggests that the selection of smaller rarefied library sizes should be approached with caution when using alpha diversity metrics, while larger normalized library sizes prevent loss of precision and reduction of the Shannon index value. However, as previously noted, the reduction in the value of the Shannon index will be dependent on the sequence variant composition of the samples.

Previous research evaluating normalization techniques has focused on beta diversity analysis and differential abundance analysis (Gloor et al., 2017; McMurdie and Holmes, 2014; Weiss et al., 2017), but the appropriateness of library size normalization techniques for alpha diversity metrics must be evaluated due to the prerequisite of having equal library sizes for accurate calculation. Utilization of unnormalized library sizes with alpha diversity metrics may generate bias due to the potential for samples with larger library sizes to inherently reflect more of the diversity in the source than a sample with a small library size. The repeated iterations of rarefying library sizes allow characterization of the variability introduced to sample diversity by rarefying at any rarefied library size (Figure 4) but does not allow evaluation of uncertainty about the diversity in the source from which the sample was taken, as is the case for all normalization-based approaches.

### 4.3 The Effects of Subsampling Approach and Normalized Library Size Selection on Beta Diversity Analysis

When samples were repeatedly rarefied to a common normalized library size with and without replacement, similar amounts of variation in the Bray-Curtis PCA ordinations were observed between the sampling approaches (Figure 5). This indicates that although rarefying with replacement seems potentially erroneous due to the fabrication of count values that are not representative of actual data, the impact on the variation introduced into the Bray-Curtis dissimilarity distances is not large and will likely not interfere with the interpretation of results. However, rarefying without replacement should be encouraged because it is more theoretically correct to represent possible data if only the smaller library size had been obtained, and it has not been comprehensively demonstrated that sampling with replacement is a valid approximation for all types of diversity analysis or library compositions.

**Figure 5.**
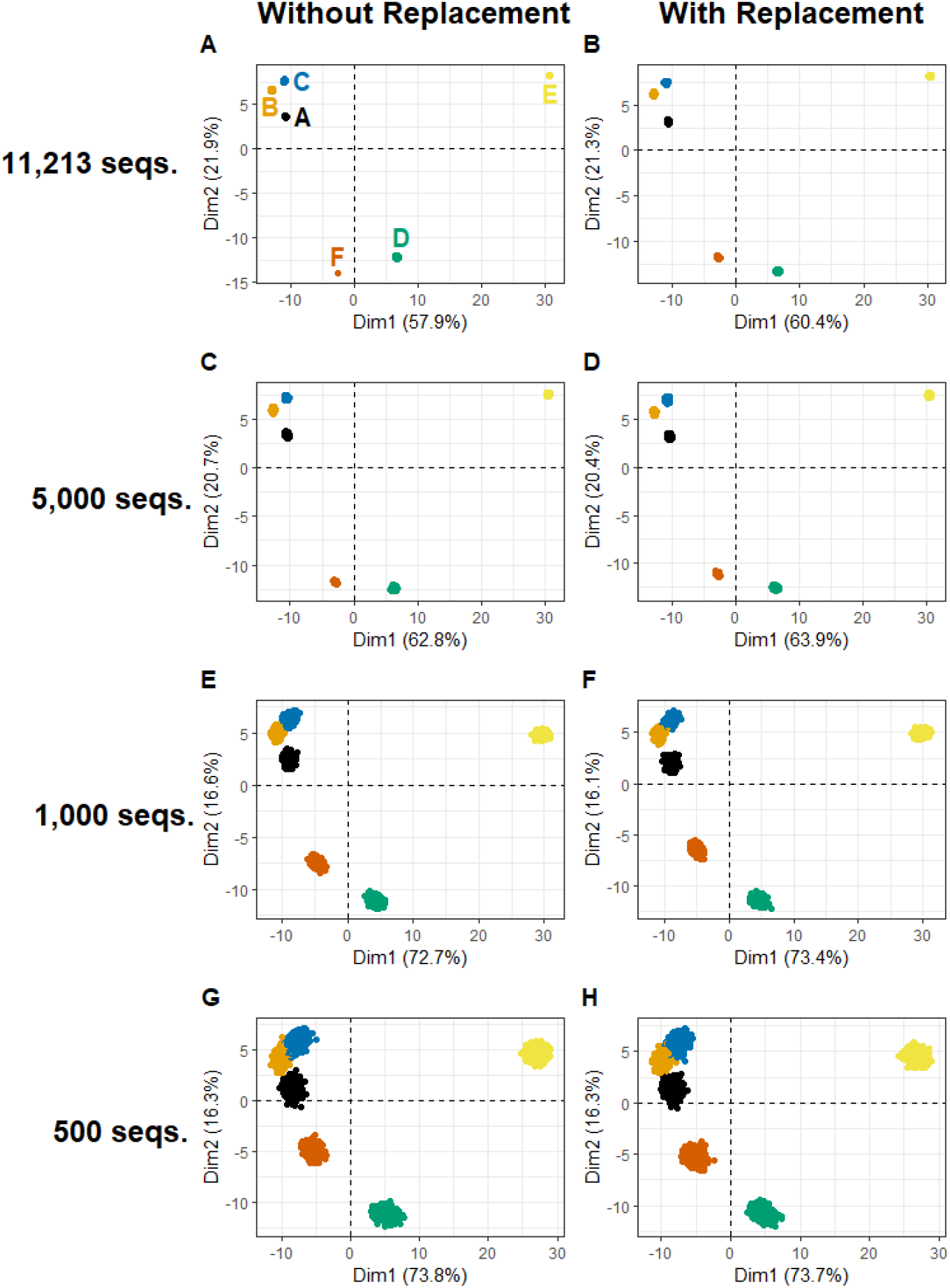
Variation in PCA ordinations (using the Bray-Curtis dissimilarity on Hellinger transformed rarefied libraries) of six microbial communities repeatedly rarefied with and without replacement to varying library sizes.

When larger normalized library sizes are maintained through rarefaction, there is less potential variation introduced into beta diversity analyses, including Bray-Curtis dissimilarity PCA ordinations. For example, in the largest normalized library size possible for these data (Figure 5A), a minimal amount of variation was observed within each community, indicating that the preservation of higher sequence counts minimizes the amount of artificial variation introduced into datasets by rarefaction (including no variation for Sample F because it is not actually rarefied in this scenario). For this reason, rarefying to the smallest library size of a set of samples is a sensible guideline. Although, a normalized library size of 5,000 is lower than the flattening portion of the rarefaction curve for samples A, B, and C (Figure 2), the selection of this potentially inappropriate normalized library size (Figure 5C) can still accurately reflect the diversity between samples without excess artificial variation introduced through rarefaction. Due to the variation introduced to the Bray-Curtis dissimilarity ordinations in the smaller rarefied library sizes (Figure 5E/G), it is critical to include computational replicates of rarefied libraries to fully characterize the introduced variation in communities. As discussed above, it has been suggested that repeatedly rarefying is inappropriate due to the introduction of “added noise”. However, as demonstrated, the maintenance of larger rarefied library sizes when repeatedly rarefying does not impact interpretation of beta-diversity analysis results. Without this replication, rarefaction to small, normalized library sizes could result in artificial similarity or dissimilarity identified between samples.

Beta diversity analysis of very small, rarefied library sizes (Figure 6A, B, C) can still reflect similar clustering patterns observed in larger library sizes but with a much lower resolution of clusters. Rarefying has previously been shown to be an appropriate normalization tool for samples with low sequence counts (e.g., <1,000 sequences per sample) by Weiss et al. (2017), which is promising for datasets containing samples with small initial library sizes or potentially analyzing subsets of data to explore diversity within specific phyla (e.g., Cyanobacteria). Caution must be taken to avoid selection of an excessively small, normalized library size due to the introduction of extreme levels of artificial variation that compromises accurate depiction of diversity (Figure 6D) and suppresses the contribution of rare variants to overall diversity. The tradeoff between rarefying to a smaller than advisable library size or excluding entire samples with small library sizes remains and can possibly be resolved by analyzing results with all samples and a small, rarefied library size as well as with some omitted samples and a larger rarefied library size.

**Figure 6.**
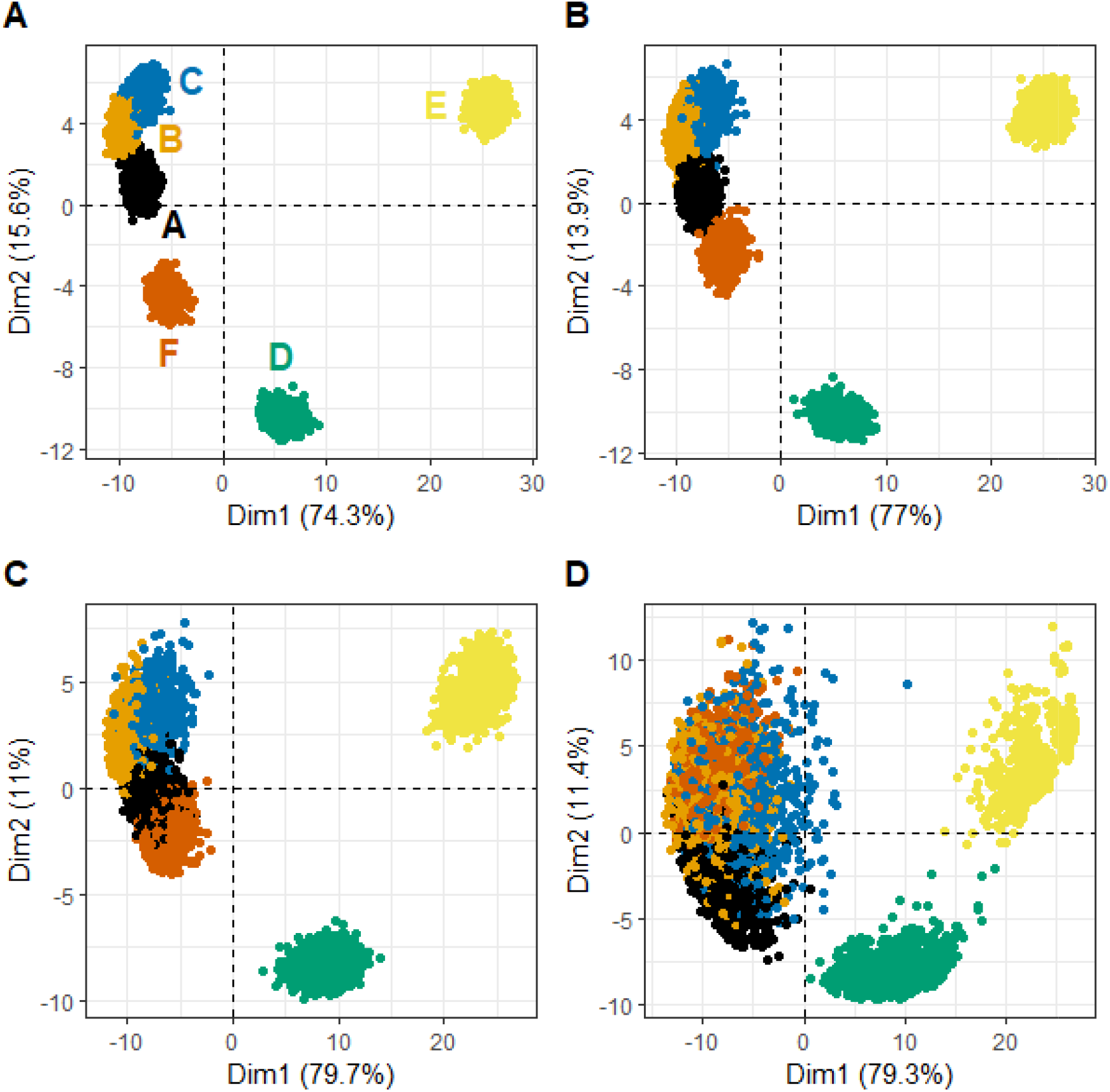
Variation in PCA ordinations (using the Bray-Curtis dissimilarity on Hellinger transformed rarefied microbial communities) of six microbial communities repeatedly rarefied to very small library sizes of (A) 400, (B) 300, (C) 200 and (D) 100 sequences.

Although rarefying has the potential to introduce artificial variation into data used in beta diversity analyses, these results suggest that rarefying repeatedly does not become problematic until normalized library sizes are very small (e.g., 500 sequences or less) for the samples considered. While we saw a degradation of the consistency and value of the alpha diversity Shannon index at 500 sequences, beta diversity analyses may be more robust to rarefaction and capable of reflecting qualitative clusters in ordination as previously discussed in Weiss et al. (2017). The artificial variation introduced to beta diversity analyses by rarefaction could lead to erroneous interpretation of results, but the implementation of multiple iterations of rarefying library sizes allows a full representation of this variation to aid in determining if apparent similarity or dissimilarity is a chance result of rarefying.

The use of non-normalized data has been shown to be more susceptible to the generation of artificial clusters in ordinations, and rarefying has been demonstrated to be an effective normalization technique for beta diversity analyses (Weiss et al., 2017). However, the use of a single iteration of rarefying does result in the omission of valid data (McMurdie and Holmes, 2014). Repeated iterations of rarefying in this study demonstrated that rarefying repeatedly does not substantially impact the output and interpretation of beta diversity analyses unless rarefying to sizes that are inadvisably small to begin with. McMurdie and Holmes (2014) were dismissive of rarefying repeatedly due to the variability it introduces, but such repetition was not evaluated in the context of beta-diversity analysis. In the case of differential abundance analysis, the added variability of rarefying would be statistically inappropriate relative to generalized linear modelling that can account for varying library sizes. Additionally, repeatedly rarefying allows for characterization of variation introduced through subsampling while accounting for discrepancies in library size, supporting the potential utility of the normalization technique for beta diversity analyses. McKnight et al. (2019) preferred use of proportions in diversity analysis over rarefying (arguing that both were superior to other normalization approaches). While proportions normalize the sum of the ASV weights to one for each sample, we note that the approach does not normalize the library size in terms of sequence counts. This is important because sample proportions will provide a more precise reflection of the true proportions of which the set of sequences is believed to be representative in samples with larger libraries than in samples with smaller libraries. In particular, using proportions of unnormalized sequence count libraries in beta diversity analysis overlooks the loss of alpha diversity associated with smaller library sizes when comparing samples with different library sizes.

### 4.4 Perspectives on Library Size Normalization

The increasing popularity and accessibility of amplicon sequencing has enabled the scientific community to gain access to a wealth of microbial community data that would otherwise not have been accessible. However, despite amplicon sequencing of taxonomic marker genes being the gold standard approach for microbial community analysis, the data handling and statistical analysis is still in the early stages of development. The diversity analyses that the scientific community desires to perform on amplicon sequencing data require library sizes to be normalized across samples, which creates the challenge of determining appropriate normalization techniques. New normalization techniques and tools are constantly being developed and released to the community with claims that the newest technique is the best and only solution that should be utilized for analysis, but they may be associated with data handling limitations, be too specifically tailored to a particular type of analysis or desired property, or not normalize the library sizes that motivated the need for normalization (McKnight et al., 2018). For example, the centered-log ratio transformation (Gloor et al., 2016) cannot be used with zero count data and amplicon sequencing datasets must be augmented with an artificial pseudocount to apply the normalization technique. The limitations of normalization techniques may affect downstream analyses, making it critical to understand the implications of the technique chosen.

Further discussion within the scientific community is needed to ensure rigorous interpretation of amplicon sequencing data without unwarranted bias introduced by the normalization technique. Approaches to microbiome data analysis that recognize data as samples from a source population and seek to draw inference about diversity in the source rather than just calculating diversity in the (transformed) sample are desirable. Random errors are inherent to sample collection, handling, processing, amplification, and sequencing and should be reflected in how resulting data are analyzed. Pending further research on such approaches, rarefying remains common in current research requiring library size normalization despite potential limitations, especially for diversity analysis. The implementation of a single iteration of rarefying is problematic due to the omission of valid data and should not be used for library size normalization. Conducting repeated iterations of rarefying, however, does not discard valid sequences and allows for the characterization of variation introduced through random subsampling in diversity analyses.

## Conclusions

- Repeated rarefying (e.g., 1000 times if computationally feasible) statistically describes possible realizations of the data if the number of sequences read had been limited to the normalized library size, thus allowing diversity analysis using samples of equal library size in a way that accounts for the data loss in rarefying.
- Rarefying with or without replacement did not substantially impact the interpretation of alpha (Shannon index) or beta (Bray-Curtis dissimilarity) diversity analyses considered in this study, but rarefying without replacement is theoretically more appropriate and will provide more accurate reflection of sample diversity.
- The use of larger normalized library sizes when rarefying minimizes the amount of artificial variation introduced into diversity analyses but may necessitate omission of samples with small library sizes (or analysis at both inclusive low library sizes and restrictive higher library sizes).
- Ordination patterns are relatively well preserved down to small, normalized library sizes with increasing variation shown by repeatedly rarefying, whereas the Shannon index is very susceptible to being impacted by small, normalized library sizes both in declining values and variability introduced through rarefaction.
- Even though repeated rarefaction can characterize the error introduced by excluding some fraction of the sequence variants, rarefying to extremely small sizes (e.g., 100 sequences) is inappropriate because the substantial introduced variation leads to an inability to differentiate between sample clusters and suppresses contribution of rare variants to diversity.
- Further development of strategies (e.g., data handling, library size normalization for diversity analyses) for ensuring rigorous interpretation of amplicon sequencing data is required.

## Acknowledgements

We acknowledge the support of the forWater NSERC network for forested drinking water source protection technologies [NETGP-494312-16]. We are also grateful for the continued support of Natural Resources Canada and Environment and Climate Change Canada in sample collection at Turkey Lakes Watershed Research Station

