## Supplemental Table 1 for "To rarefy or not to rarefy: Enhancing diversity analysis of microbial communities through next-generation sequencing and rarefying repeatedly"

Table S1: Functions from other R packages used in *mirlyn*. Mirlyn is an R package developed for library normalization and diversity analyses of amplicon sequencing and is available at [www.github.com/escamero/mirlyn](http://www.github.com/escamero/mirlyn)

| *mirlyn* function | Description | Functions used from other packages | Citations |
| --- | --- | --- | --- |
| bartax() | Generate taxonomic composition barcharts from taxonomic abundance data. | microbiome::transform(): generate compositional data from abundance data. | Leo Lahti et al.  microbiome R package.  URL: http://microbiome.github.io |
|  |  | phyloseq::tax_glom(): combine compositional data by desired taxonomic level.  ggplot2::ggplot(): plotting engine for all visualization. | Paul J. McMurdie and Susan Holmes (2013). phyloseq: An R package for reproducible interactive analysis and graphics of microbiome census data. PLoS ONE 8(4):e61217. |
| alphawhichDF() | Calculate alpha diversity values from sequence count data. | vegan::diversity(): used for alpha diversity calculation. | Jari Oksanen, F. Guillaume Blanchet, Michael Friendly, Roeland Kindt, Pierre Legendre, Dan McGlinn, Peter R. Minchin, R. B. O'Hara, Gavin L.    Simpson, Peter Solymos, M. Henry H. Stevens, Eduard Szoecs and Helene Wagner (2019). vegan: Community Ecology Package. R package version 2.5-6.    https://CRAN.R-project.org/package=vegan |
| alphacone() | Calculate alpha diversity values at different increments of library rarefaction for sequence data. | phyloseq::rarefy_even_depth(): rarefy taxonomic abundance data from multiple samples to an equal depth. | Paul J. McMurdie and Susan Holmes (2013). phyloseq: An R package for reproducible interactive analysis and graphics of microbiome census data. PLoS ONE 8(4):e61217. |
|  |  | vegan::diversity(): used for alpha diversity calculation. | Jari Oksanen, F. Guillaume Blanchet, Michael Friendly, Roeland Kindt, Pierre Legendre, Dan McGlinn, Peter R. Minchin, R. B. O'Hara, Gavin L.    Simpson, Peter Solymos, M. Henry H. Stevens, Eduard Szoecs and Helene Wagner (2019). vegan: Community Ecology Package. R package version 2.5-6.    https://CRAN.R-project.org/package=vegan |
| betamatPCA() | Principle component analysis of beta diversity values calculated from sequence count data. | vegan::vegdist(): calculate dissimilarity indices from taxonomic abundance data. | Jari Oksanen, F. Guillaume Blanchet, Michael Friendly, Roeland Kindt, Pierre Legendre, Dan McGlinn, Peter R. Minchin, R. B. O'Hara, Gavin L.    Simpson, Peter Solymos, M. Henry H. Stevens, Eduard Szoecs and Helene Wagner (2019). vegan: Community Ecology Package. R package version 2.5-6.    https://CRAN.R-project.org/package=vegan |
|  |  | vegan::decostand(): apply desired standardization to taxonomic abundance data prior to calculation of dissimilarity indices. |  |
|  |  | stats::prcomp(): principle component analysis. | R Core Team (2020). R: A language and environment for statistical computing. R Foundation for Statistical Computing, Vienna, Austria. URL    https://www.R-project.org/. |
| mirl() | Repeated rarefaction of sequence count data. | phyloseq::rarefy_even_depth(): rarefy taxonomic abundance data from multiple samples to an equal depth. | Paul J. McMurdie and Susan Holmes (2013). phyloseq: An R package for reproducible interactive analysis and graphics of microbiome census data. PLoS ONE 8(4):e61217. |
| rarefy_whole_rep() | Repeated rarefaction of taxonomic abundance data at incremental library sizes. | phyloseq::rarefy_even_depth(): rarefy taxonomic abundance data from multiple samples to an equal depth. | Paul J. McMurdie and Susan Holmes (2013). phyloseq: An R package for reproducible interactive analysis and graphics of microbiome census data. PLoS ONE 8(4):e61217. |
| rarecurve() | Visualize observed ASV count from rarefy_whole_rep() output. | ggplot2::ggplot(): plotting engine for all visualization. | H. Wickham. ggplot2: Elegant Graphics for Data Analysis. Springer-Verlag New York, 2016. |
| alphawhichVis() | Visualize alpha diversity results from alphawhichDF(). | ggplot2::ggplot(): plotting engine for all visualization. | H. Wickham. ggplot2: Elegant Graphics for Data Analysis. Springer-Verlag New York, 2016. |
| betamatPCAvis() | Plot principle component analysis from betamatPCA(). | ggplot2::ggplot(): plotting engine for all visualization. | H. Wickham. ggplot2: Elegant Graphics for Data Analysis. Springer-Verlag New York, 2016. |
|  |  | factoextra::fviz_pca_ind(): wrapper for ggplot2 visualization of PCA data. | Alboukadel Kassambara and Fabian Mundt (2020). factoextra: Extract and Visualize the Results of Multivariate Data Analyses. R package version    1.0.7. <https://CRAN.R-project.org/package=factoextra> |
| phyloseqtodf() | Creation of a dataframe from a *phyloseq* object including taxonomy, ASV read counts and metadata. |  |  |
| get_asv_table() | Generates a compiled ASV table with counts and taxonomic classification of individual amplicon sequence variants. |  |  |
| randomseqsig() | Identification of whether a taxonomic group of interest is significantly dominant or rare in the community using data shuffling. |  |  |
| plot_heat() | Generates heat maps visualizing the relative abundance of a taxonomic group of interest from a dataframe. | ggplot2::ggplot(): plotting engine for all visualization. | H. Wickham. ggplot2: Elegant Graphics for Data Analysis. Springer-Verlag New York, 2016. |
| asv_rename() | Assigns unique ASV I.D.’s to sequence variants. |  |  |
| fasta_rename() | Assigns corresponding unique ASV identifiers to a FASTA file. |  |  |
| fullbartax() | Generates taxonomic composition bar charts for specified taxonomic levels. | ggplot2::ggplot(): plotting engine for all visualization. | H. Wickham. ggplot2: Elegant Graphics for Data Analysis. Springer-Verlag New York, 2016. |
